# Ants act as olfactory bio-detectors of tumour in patient-derived xenograft mice

**DOI:** 10.1101/2022.05.16.492058

**Authors:** Baptiste Piqueret, Élodie Montaudon, Paul Devienne, Chloé Leroy, Elisabetta Marangoni, Jean-Christophe Sandoz, Patrizia d’Ettorre

**Author notes:** Equal contributions of these two authors (Jean-Christophe Sandoz, Patrizia d’Ettorre). Corresponding authors: Baptiste Piqueret and P.d.E. Current address: Max Planck Institute for Chemical Ecology, Hans-Knöll-Straße 8, 07745 Jena, Germany (Baptiste Piqueret).

## Abstract

Early detection of cancer is critical in medical sciences, as the sooner a cancer is diagnosed, the higher the chances of recovery. Tumour cells are characterized by specific volatile organic compounds (VOCs) that can be used as cancer biomarkers. Through olfactory associative learning, animals can be trained to detect these VOCs. Insects, such as ants, have a refined sense of smell and can be easily and rapidly trained with olfactory conditioning. Using urine from patient-derived xenograft mice as stimulus, we demonstrate that individual ants can learn to discriminate the odour of healthy mice from that of tumour bearing mice, and do so after only three conditioning trials. Chemical analyses confirmed that the presence of the tumour changed the urine odour, supporting the behavioural results. Our study demonstrates that ants reliably detect tumour cues in mice urine and have the potential to act as efficient and inexpensive cancer bio detectors.

## Background

With 10 million deaths, and more than 19 million cases in 2020, cancer is a leading cause of mortality worldwide (1). A way to increase cancer survival rate is to improve diagnosis methods. The sooner the tumours are detected, the higher are the chances of recovery for the patients. Current early detection methods are often invasive (e.g., coloscopy) and/or expensive (e.g., magnetic resonance imaging, MRI), therefore relatively few people profit from available detection techniques. For example, in France, the most prevalent cancer for women is breast cancer. However, despite France is a relatively rich country, less than half (48.6 % in 2019) of the population at risk (women between 50 and 74) underwent screening for this cancer.

A promising method to increase early cancer detection rate is the use of animal olfaction. Trained dogs can detect tumours in cell samples or body odour samples (2,3). This method is based on the detection of volatile organic compounds (VOCs) that are characteristic of tumours and linked to their altered cell metabolism (4). Dogs are not the only animal species used as bio-detectors of cancer. Mice were trained to discriminate tumour-bearing mice from healthy ones (5). The nematode *Caenorhabditis elegans* showed attractive chemotaxis to some cancer VOCs (6). In fruit flies, by using *in vivo* calcium imaging, a cutting edge and expensive technology, it was shown that olfactory neurons formed specific activation patterns when exposed to the volatiles from a given cancer cell line (7). Using a well-known and simple paradigm for olfactory conditioning, the proboscis extension response (8), honeybees were tested when exposed to cancer odour (9). Insects are promising detection tools as they are relatively easy to handle, do not require expensive rearing facilities, are available in large numbers and can be trained to recognize an odour in very few trials. Amongst insects, ants, and especially *Formica fusca*, demonstrated remarkable learning abilities using ecologically relevant odours. In this species, one training trial is enough to form a genuine long-term memory lasting for days. Furthermore, these ants are highly resistant to memory extinction: after training, they can be tested up to 9 times without reward before their responses start to decline (10). Recently, we tested the olfactory detection abilities of *F. fusca* towards cultured human cancer cell lines. Using ovarian (IGROV-1) and breast cell lines (MCF-7, MCF-10A and MDA-MD-231) as stimuli, we showed that ants can correctly discriminate between a cancerous cell line and a healthy one, and between two cancerous cell lines (11). Cell lines have the advantage of being easily available and to provide reproducible odour samples over time, but they do not represent the exact reality of tumours, which are complex tissues composed of different cell types. The use of tumour tissues or whole organisms is therefore a critical step for testing ants as clinical detection tools.

In this study, we used olfactory stimuli (urine samples) from patient-derived xenograft (PDX) mice carrying human tumours. Compared to cell lines that grow on a stable and known environment (culture medium), PDX mice represent a more realistic model, as the cancerous cells composing a tumour are growing in a live organism with all its complexity. Furthermore, human tumours retain their characteristics when grafted on mice (heterogeneity, relative proportion of cells, and genomic architecture) (12). Finally, tumours used in PDX are stable in time and can be duplicated, which allows for a virtually infinite number of drug tests and preclinical investigations, ensuring that the patient at the origin of the grafted tumour receives the optimal treatment (12,13).

Here, PDX were established using a “triple-negative” human breast tumour. Urine was chosen as body-odour stimuli, as it can easily be collected and stored (14). Ant workers of the species *F. fusca* were used as bio-detector since they are able to learn and detect cancer related odours stemming from standardized cancer cell lines (11). Using a simple conditioning paradigm, we tested whether ants could discriminate the urine of mice carrying a tumour from that of healthy mice. To characterize changes in VOCs composition due to the presence of cancer, urine odour samples were analysed using solid-phase micro-extraction (SPME) and gas chromatography coupled with mass spectrometry (GC-MS).

## Methods

### (a) Insects and origin of colonies

*Formica fusca* is a relatively common ant species in the Northern hemisphere. Colonies may contain several hundred individuals and are headed by one queen (monogynous) or several queens (polygynous). Three queenright colonies were collected in the forest of Ermenonville (France, 49°09′51.5″ N, 2°36′49.2″ E) and kept under laboratory conditions (25 ± 2 °C, 50 ± 10% relative humidity, 12 h/12 h: day/night) at the Laboratory of Experimental and Comparative Ethology (LEEC, University Sorbonne Paris Nord). Tested ants were foragers (ants that leave the nest to search for food) and were individually marked with a dot of oil-based paint (Mitsubishi Pencil) on the abdomen and / or thorax one day prior to the conditioning. In all cases, each individual ant was used only for one conditioning and test experiment.

### (b) Mice and odour source

Female immunodeficient mice (Swiss nude mice from Charles River Laboratories) were maintained under specific pathogen-free conditions at the Laboratory of pre-clinical investigation (LIP, Institut Curie, Paris, France). The housing facility was kept at 22°C (± 2°C) with a relative humidity of 30-70%. The light/dark cycle was 12 h light/12 h dark. Patient-derived xenografts (PDX) from a triple-negative breast cancer (HBCx-11, mesenchymal subtype (15)) were established with consent of the patient. The study was approved by the local ethics committee (Breast Group of *Institut Curie* Hospital). Twelve mice were used in our study: half (n = 6) were graft with tumour at 9-week-old into their interscapular fat pad, the other half (n = 6) underwent the same chirurgical procedure but without the graft of the tumour (healthy sham group). Mice were kept for seven weeks and were euthanized at the end of the experiment before the PDX had a critical size of 15/15 mm. Tumour size was measured with a manual calliper, and the total volume of a tumour was calculated as *V* = *a* x *b*^2^ / 2, *a* being the largest diameter, *b* the smallest (Table 1 gives details about the size of each tumour). Care and housing of animals were in accordance with Institutional Animal Care and French Committee–approved criteria (project authorization no. 02163.02).

**Table 1.** Size of the tumour of each mouse 7 weeks after the initial graft. The estimated volume of tumour (V) was calculated as V = *a* x *b*^2^ / 2, with *a* being the largest side of the tumour and *b* the smallest side of the tumour.

| Identity of the mouse | Largest side of the tumour (mm) | Smallest side of the tumour (mm) | Estimated volume (mm<sup>3</sup>) |
| --- | --- | --- | --- |
| #1 – 6 (Sham) |  |  |  |
| #11 | 10 | 9 | 405 |
| #12 | 12 | 11 | 726 |
| #13 | 11 | 11 | 665.5 |
| #14 | 13 | 11 | 786.5 |
| #15 | 9 | 7 | 196 |
| #16 | 11 | 10 | 550 |
| Mean size | 11 | 9.83 | 554.83 |

Urine of mice was used as olfactory stimuli and was collected seven weeks after the graft. Mice were placed individually in a cage (43 cm x 27 cm, height = 16 cm) without food and water for 3h. The floor of the cage was covered with filter paper, which was collected at the end of the 3h soaked with urine. Samples of 1 cm^2^ of urine imbibed filter paper were inserted individually in an Eppendorf tube and stored at −20 °C.

### (c) Experimental protocol

The experimental protocol was similar to the one used in Piqueret et al., (10). In brief, we trained individual ants to associate an odour (Conditioned Stimulus - CS, urine of mice) with a reward (Unconditioned Stimulus - US, 30% sugar solution) using a circular arena (Figure 1A). Three conditioning trials were performed by each ant (Figure 2). The time needed by the ant to find the reward was the variable measured in each conditioning trial.

**Figure 1.**
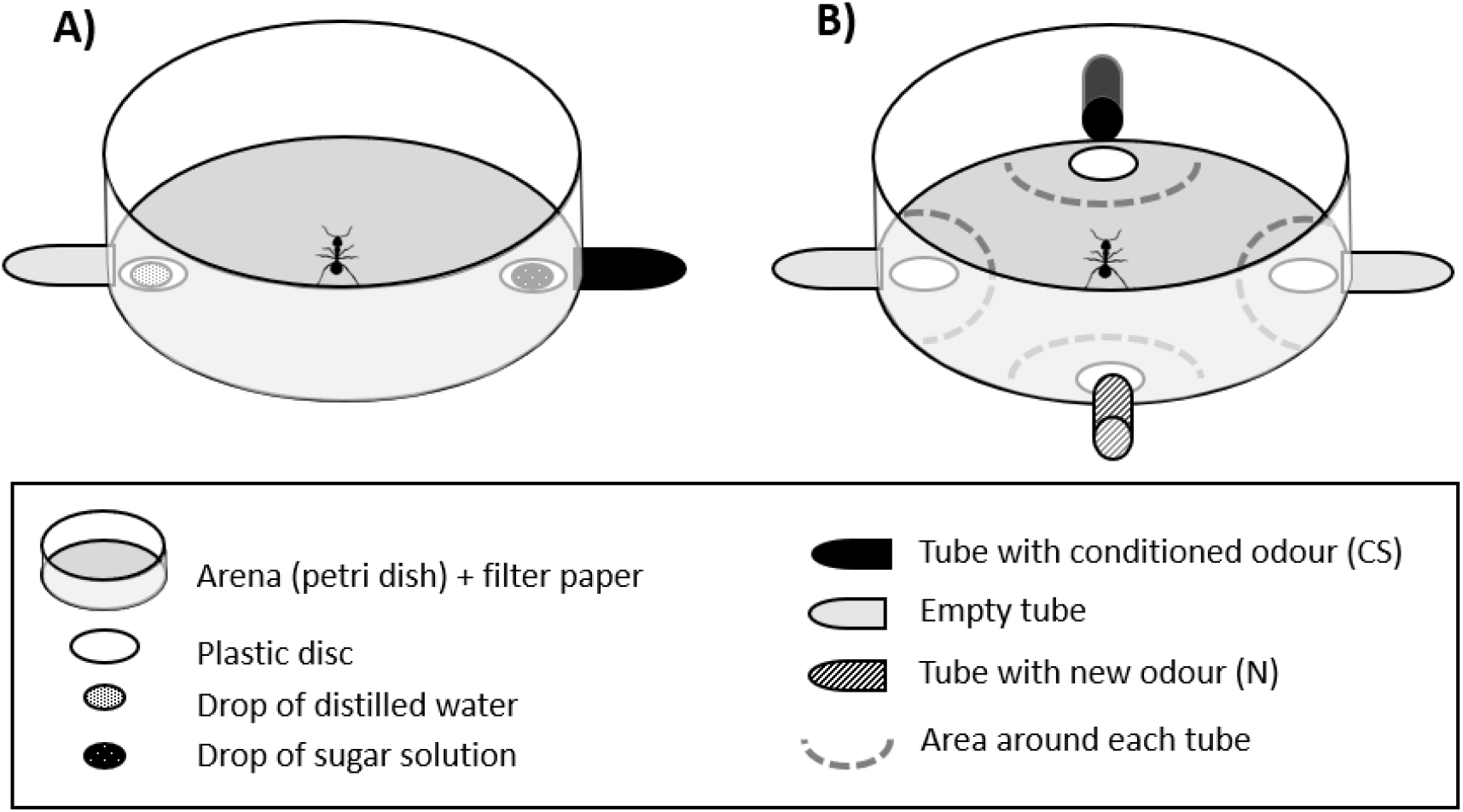
Schema of the experimental device (arenas). A: Set-up used during the conditioning phase. The time to find the reward was noted. B: Slightly modified set-up used during the memory tests. The time spent by the ant in the vicinity of the odours (areas, in dashed lines) was recorded during two minutes. Each area measured 35.5 cm^2^. The orientation of the arena in the experimental room was changed between trials so that ants could not learn visual cues.

**Figure 2.**
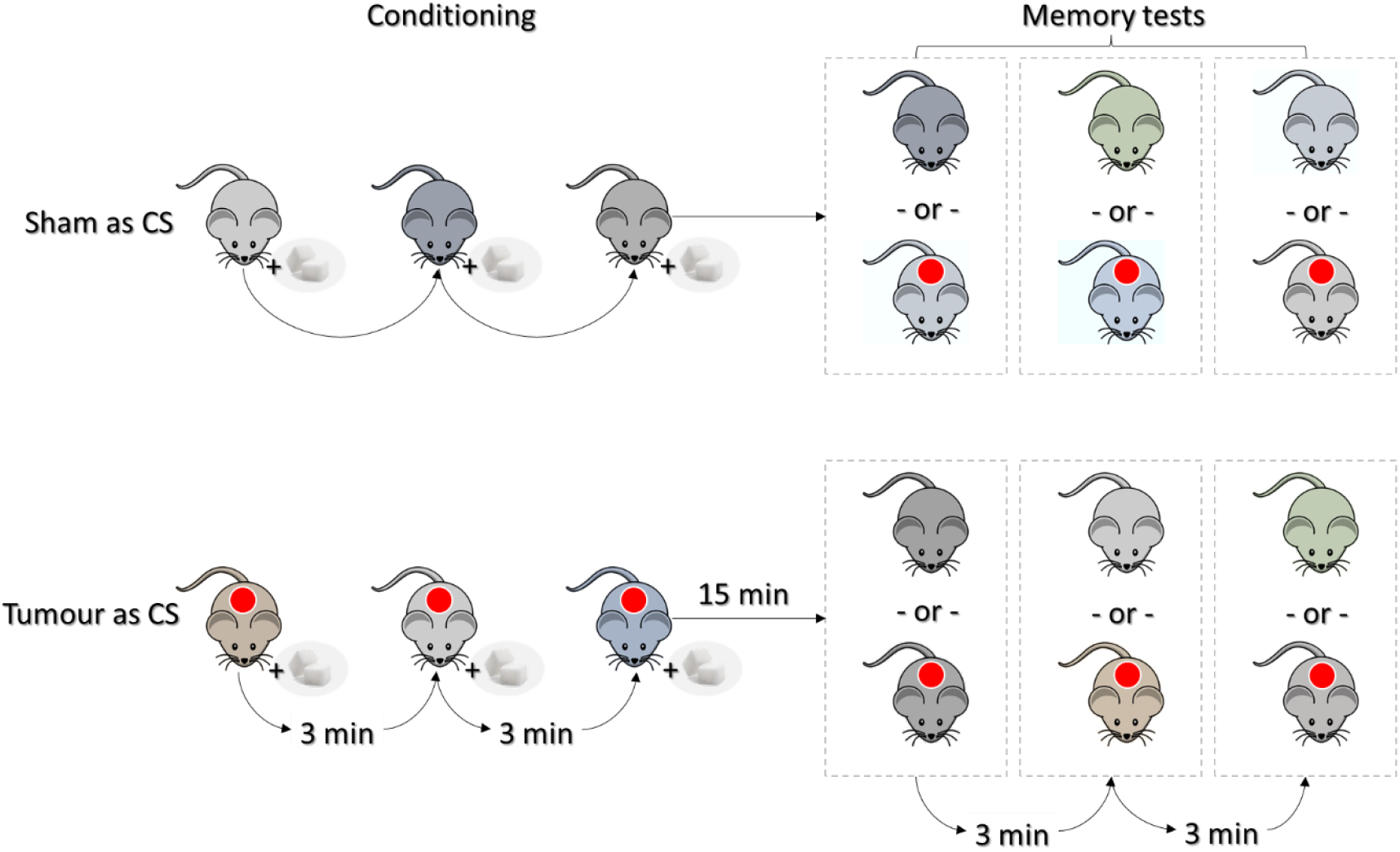
Schema of the procedure ants underwent during the behavioural experiments. Individual ants were trained to associate the odour (urine) of mice to the reward (sugar solution) three times during the conditioning procedure, and later tested with odours from other mice during the memory tests. Ants were trained either with urine of mice from the *sham* group (sham as CS, above), or from the *tumour* group (tumour as CS, below). The red circles indicate that mice have a tumour. During the memory tests, ants had to choose between urine of individuals from the two groups (*tumour* vs *sham*). Each ant encountered urine samples from 9 different mice during the full procedure. They never encountered the same stimulus more than once to exclude the possible use of individual identity cues.

To test if ants had learned the CS, we performed memory tests without reward using an arena similar to the one used for the conditioning (Figure 1B). In these memory tests, two odorant stimuli were presented at the same time, on each side of the arena. One was the CS, while the other one was a new stimulus (N). When urine from a mouse with tumour was used as CS, we used urine from a sham-mouse as N, and vice versa. Two unscented stimuli were also present (empty Eppendorf tube) to control for the potential learning of visual cues. Circular areas were drawn around each stimulus, which allowed us to record as variable the time spent by the ant in the vicinity of each odorant. Each memory test lasted 2 min and the behaviour of the ant was scored using a behavioural transcription tool (software Ethoc v. 1.2, CRCA, Toulouse, France). All experiments were video recorded with a camera (Canon, Legria HFR806) placed above the experimental arena. Ants underwent three consecutive unrewarded memory tests at 15, 20, and 25 min after the end of the last conditioning trial (Figure 2). We used different stimuli (urine from different mice) in the conditioning trials and in the memory test. Therefore, ants could not rely on the possible individual odour of a given mice between the training and the tests.

The same stimuli were used for a group of six ants, which is one advantage of using urine stimuli (14). However, we tested whether significant differences in terms of memory abilities were present when ants were tested with *fresh* samples (urine never used in a test) or *reused* one (used for at least another test). Ants tested with *reused* samples did not perform differently from ants tested with *fresh* samples (*stimulus* × *sequence*: F = 1.20, d.f = 2, p ≥ 0.1).

### (d) Chemical analysis

Odour emitted by the urine of mice used in the behavioural experiments were characterized using chemical analysis. A filter paper (1 cm^2^) soaked with urine was inserted in a 15 mL glass vial and placed at 37 °C using a water bath. A SPME fibre (50/30 DVB/CAR/PDMS, Supelco) was introduced through the PTFE/silicone 1.5 mm cap of the vial for 50 min (11,16). After that, the fibre was immediately inserted into an Agilent Technologies 7890A gas-chromatograph, equipped with a HP5MS GC column (30 m × 0.25 mm × 0.25 μm, Agilent Technologies, Les Ulis Cedex, France). The carrier gas was helium (1 mL.min^−1^), and the injection was split less (250 °C). The oven temperature was programmed at 40 °C for 5 min, then increased to 220 °C at 7 °C.min^−1^, and to 300 °C at 15 °C.min^−1^ and was held for 3 min. The GC was coupled with a 5975 C mass-spectrometer (Agilent Technologies). Mass spectra were recorded with electron impact ionization at 70 eV. Peak areas were integrated with MSD ChemStation software version E.02.01.1177 (Agilent Technologies). Peaks were identified by comparing their ion spectrum to the NIST library (NIST v2.2, 2014) and to standards injected with the same temperature program (benzyl alcohol, acetophenone, nonanal, dodecane, decanal, all from Sigma Aldrich, Saint-Louis, MO, USA).

### (e) Statistics

#### (a) Behaviour analysis

Data were analysed using R software (v. 4.0.0, (17)). Significance was fixed at α = 5%. All data were analysed using linear mixed models (LMMs, *lmer* function from the package *lme4*, (18)). Normality of dataset was obtained either with log function (for the conditioning dataset) or with square root transformation (for the memory tests dataset). The identity of the ants and the colony were included as nested random factors.

##### a. Acquisition

We analysed the effect of the number of conditioning trials (factor *trials*) on the dependent variable *time* (continuous variable, the time to find the reward). We analysed the effect of the *conditioning odorant* (factor with two levels, tumour bearing mice or sham mice). We also looked at the interaction *conditioning odorant* × *trials* to detect possible differences in ants’ responses depending on the odorant used.

##### b. Memory tests

First, we checked whether ants spent more time near the tubes with odours than near the control unscented tubes by analysing the effect of the independent variable *presence of odour* (factor with two levels, yes or no) on the dependent variable *time* (continuous variable, the time spent near the odorant vials or the unscented ones). Then, in all experiments, we analysed the effect of the independent variable *stimulus* (factor with two levels, CS or N) on the dependent variable *time* (continuous variable, the time spent in the vicinity of a stimulus) during the 3 memory tests. Finally, we analysed the memory tests separately for each stimulus origin (*tumour* or *sham* stimulus).

### 2. Chemical analysis

The areas of 49 regularly occurring peaks were standardized by calculating the ln (Pi /g(P)) (19), where Pi is the area of a peak and g(P) is the geometric mean of all the peak areas of the individual. On these variables, we run a Principal Component Analysis (PCA) (Table S3) and compared the scores of the first two principal components (PC1, explaining 45.57% of the total variance, and PC2 that account for 29.90%) between groups (urine of tumour mice vs sham mice) using Wilcoxon’s rank sum test.

We then calculated the chemical distance of each tumour sample from the centroid of the sham group and we tested whether this distance could be correlated to the size of the tumour.

## Results

### (a) Ants can associate the odour mice urine to a food reward

We conditioned half of the ants with the urine of tumour-bearing mice and the other half with urine of heathy, sham treated mice. During the acquisition phase, the ants learned the two different stimuli (urine from tumour mice and from sham mice) in a similar way, as shown by the absence of any significant *conditioning odorant* × *trial* interaction (p > 0.1). The time spent by the ants to find the reward decreased significantly across the three training trials (p < 0.001, Table S1). This suggests that ants learned that the urine odour (CS) predicted the sucrose reward (Figure 3A). Memory tests without reward were then performed to assess if ants could recognize the learned urine type (tumour or healthy).

**Figure 3.**
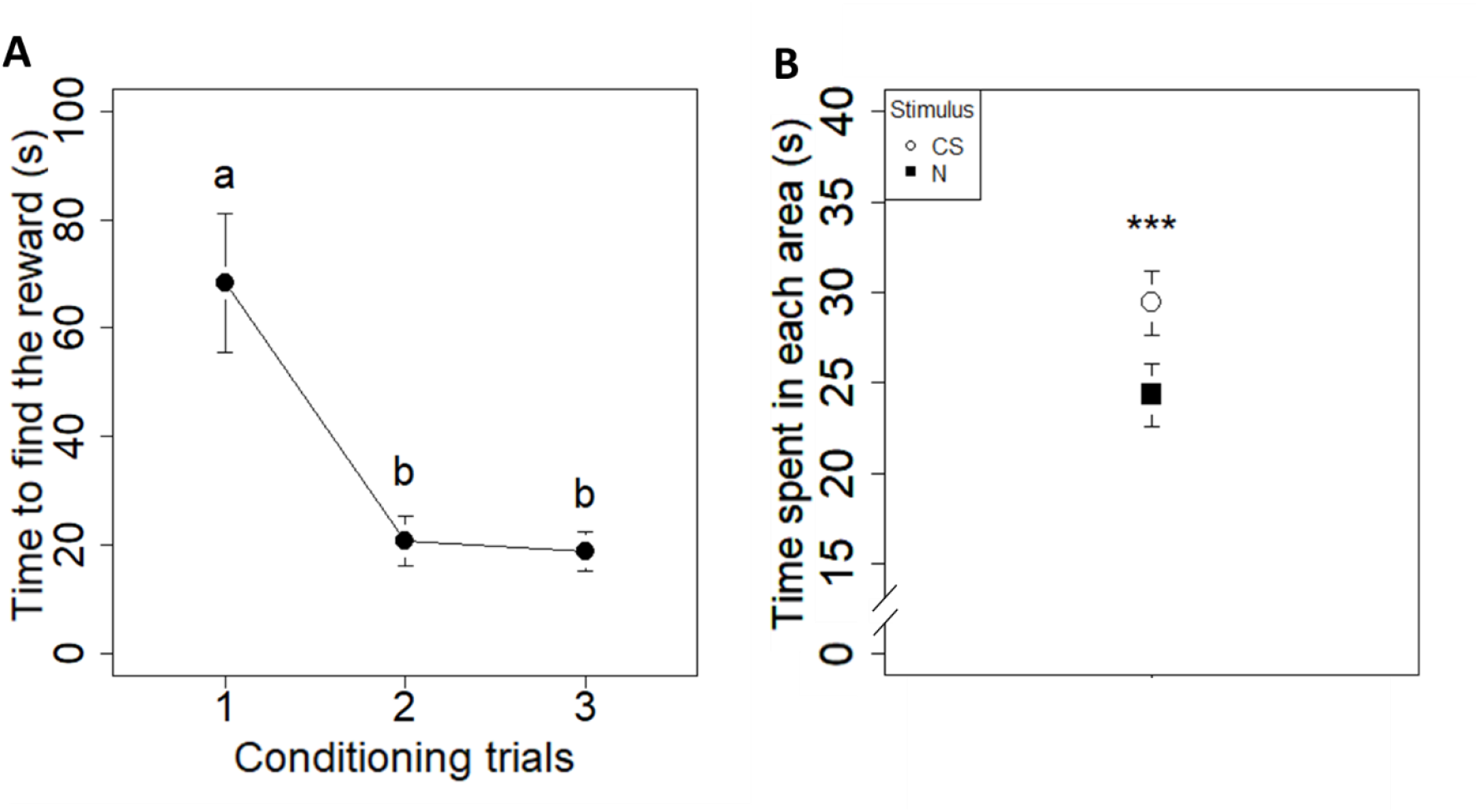
Ants’ performances during acquisition and memory tests in an appetitive conditioning experiment with urine of mice as odour stimulus. A: Ants were conditioned to associate the odour of mice urine (N = 70) either from the tumour group (N = 35) or from the sham group (N = 35) with a reward (sugar solution 30%). The graph shows the time needed by the ants to find the reward in the circular arena (Figure 1). Three conditioning trials were done with an inter-trial interval of three minutes. Significant differences between conditioning trials are indicated with different letters. B: time spent by individual ants in the vicinity of each stimulus during the memory tests (circle: conditioned stimulus, CS area; square: novel odour, N area). Circles and squares represent the mean while error bars show confidence intervals (95%). Significant differences between stimuli are noted with asterisks (***: p ≤ 0.001).

### (b) Ants can differentiate tumour-bearing mice from heathy ones after a rapid learning phase

Ants spent more time near the odorant tubes than near the unscented ones (p < 0.001), indicating that ants were not conditioned to the presence of a tube (visual cue), but to the odour. When focusing on the time spent near the tubes with odours, ants conditioned with the urine of tumour mice as CS and ants conditioned with the urine of healthy mice as CS behave similarly, as shown by the absence of significant *stimulus* × *conditioning odorant* interaction (p > 0.1). During the memory tests, ants spent more time near the CS odour than near the N odour independently of the odour used during conditioning (p < 0.001, Figure 3B). Analysing the two different subsets of data separately gives similar results (ants conditioned with tumour odour: p = 0.001; ants conditioned with healthy odour: p = 0.006, Table S2).

These results indicate that ants are able to detect whether mice were sick (presence of a tumour) or healthy based solely on the odour of their urine.

### (c) Discrimination based on urine VOCs

To investigate the cues used by ants to differentiate the tumour mice from the healthy ones, we analyzed the urine composition with Solid-Phase Micro-Extraction (SPME) coupled with Gas Chromatography and Mass Spectrometry (GC-MS). The two groups of mice (tumour-bearing and healthy) can be discriminated using PCA. Six principal components (PCs) with eigenvalues > 1, explaining together more than 90% of the total variance (Table S3) were used for discrimination. The first two PCs accounted for 75.46% of the total variance (Table S3). The two groups (*Tumour* and *Sham* group) are statistically different for the scores of the PC2 (p < 0.05), but not for PC1 (p > 0.1). Indeed, in the plot of the first two PCs the confidence ellipses are partially overlapping each other (Figure S1). In particular, the samples from two *tumour* individuals (#11 and #15) are similar to those from the sham group. We thus investigated whether this could be linked to the size of the tumour.

In fact, mice #11 and #15 were the individuals with the smallest tumours (respectively 405mm^3 &^ 196 mm^3^, Table 1), well below the mean tumour size (554.83mm^3^). We then reanalysed the data considering a group of mice with *small* tumour (below the mean size), and a group with *big* tumours (above the mean size, Table 1). With these new categories, the samples of mice with the big tumours are very well separated from those of the sham individuals (Figure 4A, PC2: p < 0.01).

**Figure 4:**
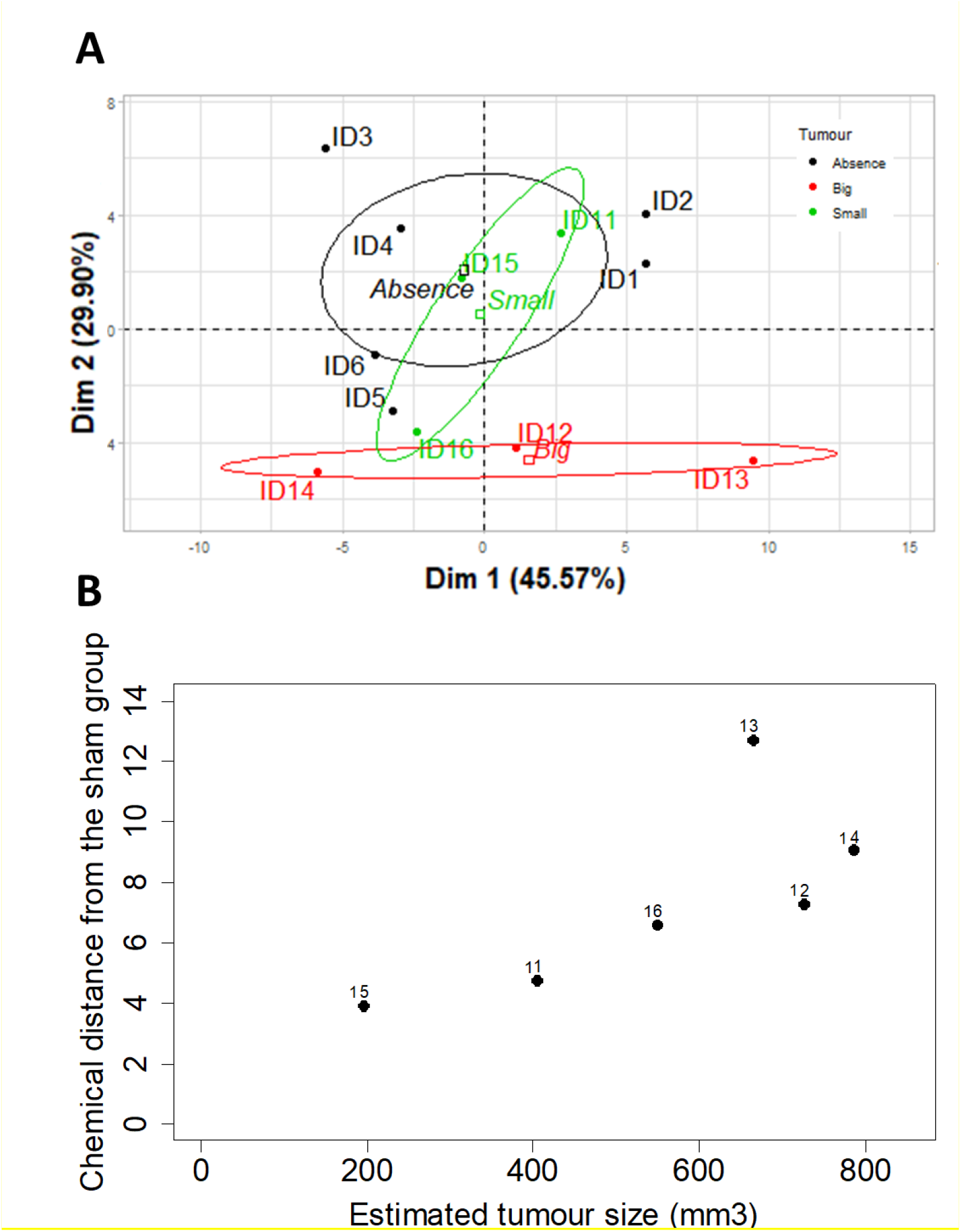
Chemical differences induced by the tumour presence in mice. **A**: Visualization of the categorization of the urine samples using the first two Principal Components (PC), explaining 68.82% of the total variance. Ellipses represent the confidence interval (0.95). The individuals with large tumour (red) are very well separated from the sham individuals (black). The ones with small tumour (green) are between the two other groups. **B**: Relationship between tumour size of mice and chemical distance of their urine samples from those of the sham group. Estimated tumour sizes (x-axis) are plotted against the chemical distance of each tumour sample from the centroid of the sham group (y-axis). Data are highly correlated (Pearson correlation test, rp = 0.73, p < 0.1).

Using the first 6 PCs (> 90% of the total variance, Table S3), we calculated the chemical distance (Euclidean distance) of each sample of tumour bearing mice from the mean coordinates (centroid) of the samples of sham group (Figure 4B). We found a strong link between the size of the tumour and chemical distance from the sham group: the larger the size of tumour, the greater the distance from the sham group (r = 0.73, p < 0.1), although the p-value shows only a tendency given to the sample size (n=6). Identification of the detected peaks can be seen in Table S4.

## Discussion

We demonstrate that ants are able to learn to associate a complex blend of odours with a reward, after only three conditioning trials. This simple and fast conditioning protocol allowed *Formica fusca* ants to discriminate tumour-bearing PDX mice from tumour-free PDX mice in unrewarded tests. This discrimination is based on the odour of the urine of mice altered by the presence of a tumour. Using chemical analysis, we also demonstrate that the larger a tumour is, the more different the odour of the bearing individual is from that of the control.

*F. fusca* ants are fast learners and show impressive memory abilities after only three associations between an odour and a reward. Such learning abilities were already described in this species (10), but also in other ant species, like *Lasius niger* (20), *Camponotus spp*. (21–23) and *Linepithema humile* (24,25). Odours used in these studies were pure compounds (e.g. hexanal) or odour blends (e.g. food flavouring), which were always ecologically relevant for ants, as they were based on floral or fruit odours. In our previous study focused on learning of cancer-related odours by ants (11), *F. fusca* ants were able to associate the odour of cultured human cancer cells (IGROV-1, MCF-7, MCF-10A, or MDA-MD-231) with a reward. Obviously, cancer cells are not ecologically relevant for ants, but many animals are known to be able to learn olfactory stimuli that are not present in their natural environment, such as drugs or explosives, given the right incentive (26). In our previous study on cancer odours, we showed that ants are able to discriminate a cancerous cell line from a healthy one, and a cancerous cell line from a different cancerous cell line.

PDX mice established with human breast tumours are a step forward in clinical cancer detection and represent an even more complex task for bio-detection as *i)* the odour used as stimulus is not directly extracted from the cancer cells, *ii)* tumours are not composed of a single cell type and are therefore heterogeneous, and *iii)* cancer cell odours might be overshadowed by the body odours of individuals or altered when passing from the local micro-environment of the tumour tissue to the body fluids used as conditioning stimulus (e.g. blood, exhaled breath, faeces, sweat, or urine). Despite all this complexity, ants reliably discriminated tumour-free urine samples (*sham* group) from tumour-carrying ones (tumour group) independently of the odour used during conditioning.

Our chemical analysis based on the volatiles emitted by urine, showed that odour samples from the tumour group are chemically different from those of the sham group, and that this difference is correlated to the size of the tumour (the larger the tumour, the more dissimilar the odour of the mouse is from the control). These results are in accordance with the behaviour results, in which ants discriminated the tumour-carrying and tumour-free mice based on olfaction.

In this study, we used urine from mice with a well-developed tumour (few days before the ethical sacrifice) as odour stimuli. We noted that the level of alteration of the mice odour was linked to the development of the tumour. Based on our chemical analysis, we found that some tumour-bearing individuals were similar to the sham ones. This does not mean that ants were not able to discriminate the small tumour individuals from the sham ones, as ants are able to detect compounds at a very low concentrations (27). During our behaviour tests, we randomly assigned each ant to the odour of three different mice during the conditioning, and six other mice during the memory tests (three from the conditioned group, three from the other group). In this way, ants could not rely on the possible individual odour of a given mouse. However, this experimental plan was not designed to quantify the impact of a weak stimulus (a small tumour) on the ant performances. It would be interesting to test the olfactory abilities of ants using only small and similar sized tumours, in order to find their detection threshold. From a clinical point of view, the optimal conditioning protocol would have the shorter delay possible between the screening and the result. Using cancer related odour (11), we managed to pin down the conditioning time of ants to 30 minutes (three conditioning trials). *F. fusca* ants have the ability to learn a simple odour stimulus after a single presentation (10), and we aim at testing this ability using complex cancer-related odours. Finally, would it be possible to train not a single ant at a time, but hundreds of individuals at the same time? Social insects (such as ants) live in colonies, and communication is essential for them (28). It was demonstrated in another social insect species that one trained honeybee could transfer a learned association to a naïve bee when their antennae were in contact (29). Studying how information are shared amongst nestmates could provide an exponentially faster conditioning methods if cancer related odours are shown to be successfully shared within the colony.

In this first study combining olfactory abilities of ants and the use of body fluids from human tumours in PDX mice, we found that ants can be used as bio-detectors, as they reliably discriminate healthy individuals from tumour-bearing ones, and vice versa. Ants could thus become a fast, efficient, inexpensive, and non-invasive tool for detection of human tumours.

## Author’ Contributions

B.P.: Conceptualization, Data curation, Formal analysis, Investigation, Methodology, Software, Visualization, Validation, Writing – original draft, Writing – review & editing. E.Mo.: Resources, Writing – review & editing. P.D: Methodology. C.L: Investigation, Formal analysis, Validation. E.Ma: Resources, Writing – review & editing. J-.C.S.: Conceptualization, Formal analysis, Funding acquisition, Project administration, Supervision, Validation, Writing – review & editing. P.d.E: Conceptualization, Formal analysis, Funding acquisition, Project administration, Supervision, Validation, Writing – review & editing

## Acknowledgments

We are grateful to Heiko Rödel for statistical assistance with behavioural analysis. We thank Fatima Mechta-Grigoriou for making this collaboration between the LEEC and the LIP possible.

