## Supplementary material for "Ants act as olfactory bio-detectors of tumour in patient-derived xenograft mice": Figure S1 & Tables S1-S4

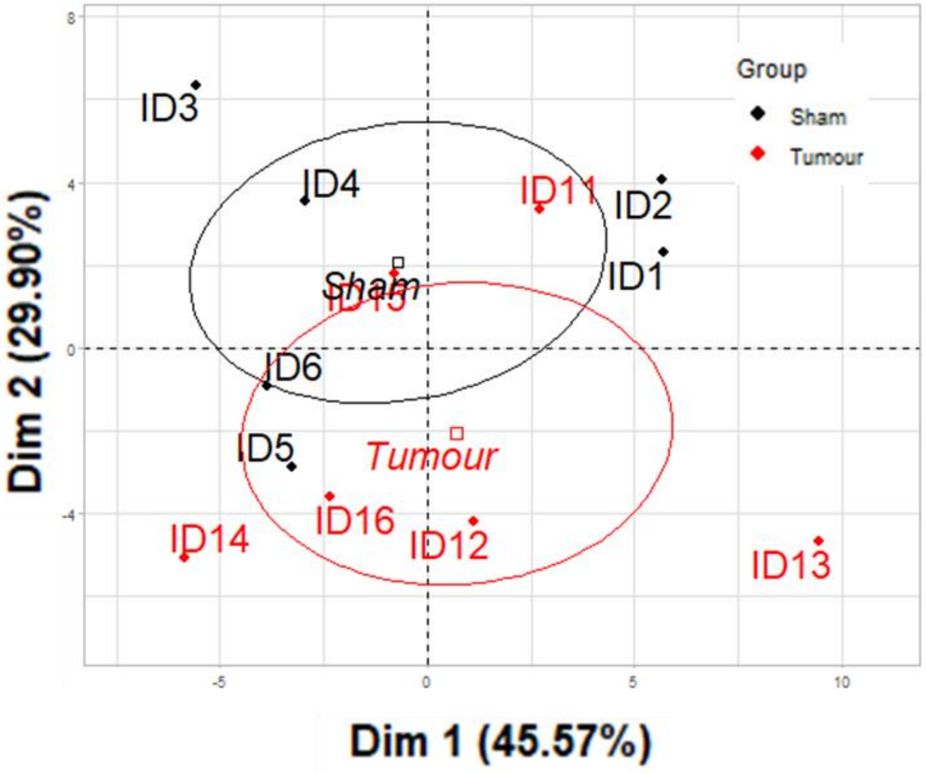

**Fig. S1. First PCA with the two original groups.** Visualizations of the categorization of the urine samples using the first two Principal Components (PC), explaining 68.82% of the total variance. Ellipses represent the confidence interval (0.95). The two groups are partially overlapping each other. Two individuals in particular from the tumor group are within the sham group. The mouse 11 and the 15. Both have the smallest tumors.

**Table S1. Result of the different models used for analyzing ants' conditioning performances.** Time was always the dependent variable. Significant effects ($p < 0.05$) are indicated in bold.

| Focus on the factor(s) | Df | F value | p value |
|---|---|---|---|
| Tumour factor experiment | | | |
| Conditioning odorant × trials | 2 | 0.2899 | 0.7488 |
| Conditioning odorant | 1 | 0.6879 | 0.4098 |
| Trials | 2 | 59.7795 | **<0.001** |
| Trials (1 *vs* 2) | 1 | | **<0.001** |
| Trials (1 *vs* 3) | 1 | | **<0.001** |
| Trials (2 *vs* 3) | 1 | | 1 |

**Table S2. Results of the model used for analysing ant's performances in the memory tests.** Time was the dependent variable. Significant effects ($p < 0.05$) are indicated in bold. Tendency effects ($p < 0.1$) are underlined.

| Focus on the factor(s) | Df | F value | p value |
|---|---|---|---|
| Global effects | | | |
| Presence of odour | 1 | 67.4 | **<0.001** |
| Stimulus × Trial × conditioning odorant | 1 | 0.3183 | 0.5729 |
| Trial × Conditioning odorant | 1 | 0.0144 | 0.9047 |
| Stimulus × Trial | 1 | 0.0376 | 0.8463 |
| Stimulus × Conditioning odorant | 1 | 0.0882 | 0.7666 |
| Trial | 1 | 0.2495 | 0.6177 |
| Stimulus | 1 | 18.0584 | **<0.001** |
| Stimulus (memory test 1) | 1 | 13.312 | **<0.001** |
| Stimulus (memory test 2) | 1 | 1.6498 | 0.2011 |
| Stimulus (memory test 3) | 1 | 6.9994 | **0.0091** |
| Tumour odour as CS | | | |
| Stimulus × Trial | 1 | 0.3029 | 0.5826 |
| Trial | 1 | 0.0756 | 0.7836 |
| Stimulus | 1 | 10.8299 | **0.0012** |
| Stimulus (memory test 1) | 1 | 10.407 | **0.0019** |
| Stimulus (memory test 2) | 1 | 0.4074 | 0.5254 |
| Stimulus (memory test 3) | 1 | 4.9939 | **0.0287** |
| Sham / Healthy odour as CS | | | |
| Stimulus × Trial | 1 | 0.0659 | 0.7976 |
| Trial | 1 | 0.1841 | 0.6683 |
| Stimulus | 1 | 0.1841 | **0.0069** |
| Stimulus (memory test 1) | 1 | 3.6382 | <u>0.0607</u> |
| Stimulus (memory test 2) | 1 | 1.5843 | 0.2126 |
| Stimulus (memory test 3) | 1 | 2.6185 | 0.1103 |

**Table S3. Results of a Principal Components analysis (PCA).** Results are based on the normalized peak area of 49 peaks extracted from the chemical analysis of the urine samples. We selected 6 principal components (PCs), which together explain more than 90% of the total variance.

| | PC1 | PC2 | PC3 | PC4 | PC5 | PC6 |
|---|---|---|---|---|---|---|
| **Eigen value** | 22.33 | 14.65 | 3.57 | 2.39 | 1.84 | 1.24 |
| **Variance** | 45.57 | 29.90 | 7.29 | 4.89 | 3.77 | 2.53 |
| **Cumulative Variance** | 45.57 | 75.46 | 82.75 | 87.64 | 91.40 | 93.94 |

**Table S4. VOCs found in samples, with their chemical identity when available.** VOCs in bold were confirmed with injected standards using the same protocol. The spectra of other VOCs were compared with a reference database (NIST v2.2. 2014). HC means that the compound is a hydrocarbon. Few compounds were not identified or categorized as the percentage of concordance with the database was too low.

| Peak No. | Compounds |
|---|---|
| **1** | 2-Hexenal, 2-ethyl- |
| **2** | 2-Heptanone |
| **3** | Dimethyl sulfone |
| **4** | 3-Heptanone, 6-methyl- |
| **5** | 6-Hepten-3-one, 4-methyl- |
| **6** | 2-Ethyl-1-hexanol (33%)+**Benzyl alcohol** |
| **7** | 7-Exo-ethyl-5-methyl-6,8-dioxabicyclo[3.2.1]oct-3-ene |
| **8** | 4,6-Dimethyldecane (4,6diMeC10) |
| **9** | **Acetophenone** |
| **10** | **Nonanal** |
| **11** | Benzyl methyl ketone |
| **12** | **Dodecane** |
| **13** | **Decanal** |
| **14** | Dodecane, 2,6,11-trimethyl- (21%) (2,6,11 triMeC12) |
| **15** | HC |
| **16** | Tridecane |
| **17** | HC |
| **18** | 4,8-Dimethyldodecane (4,8diMeC12) |
| **19** | -------------- |
| **20** | HC |
| **21** | HC |
| **22** | -------------- |
| **23** | -------------- |
| **24** | Tridecane, 4-methyl (4MeC13) |
| **25** | Propanoic acid, 2-methyl-, 3-hydroxy-2,2,4-trimethylpentyl ester |
| **26** | 3,5-Dibutoxy-1,1,1,7,7,7-hexamethyl-3,5-bis(trimethylsiloxy)tetrasiloxane |
| **27** | Tetradecane (22%) |
| **28** | Tetradecane, 7-methyl- (7MeC14) |
| **29** | HC |
| **30** | -------------- |
| **31** | 6MeC15 |
| **32** | HC |
| **33** | di MeC14 |
| **34** | 2,6,10-Trimethyltridecane (28%) (2,6,10 triMeC13) |

| | |
|---|---|
| **35** | Pentadecane, 5-methyl- (5MeC15) |
| **36** | HC |
| **37** | -------------- |
| **38** | HC |
| **39** | x,9 diMeC15 |
| **40** | HC |
| **41** | HC |
| **42** | Butylated Hydroxytoluene |
| **43** | -------------- |
| **44** | HC |
| **45** | Tetradecane, 2,6,11-trimethyl- (2,6,11triMeC14) |
| **46** | HC |
| **47** | HC |
| **48** | HC |
| **49** | HC |
